# Brefeldin A and M-COPA block the export of RTKs from the endoplasmic reticulum via simultaneous inactivation of ARF1, ARF4, and ARF5

**DOI:** 10.1101/2023.10.11.558504

**Authors:** Miyuki Natsume, Mariko Niwa, Sho Ichikawa, Takuma Okamoto, Hisazumi Tsutsui, Daiki Usukura, Takatsugu Murata, Ryo Abe, Motoyuki Shimonaka, Toshirou Nishida, Isamu Shiina, Yuuki Obata

**Affiliations:** Laboratory of Intracellular Traffic & Oncology, National Cancer Center Research Institute, Tsukiji, Chuo-ku, Tokyo 104-0045, Japan; Department of Applied Chemistry; Department of Chemistry, Faculty of Science, Tokyo University of Science, Shinjuku-ku, Tokyo 162-8601, Japan; Tokyo University of Science, Noda Chiba 278-8510, Japan; National Cancer Center Hospital, Tsukiji, Chuo-ku, Tokyo 104-0045, Japan; Laboratory of Nuclear Transport Dynamics, National Institutes of Biomedical Innovation, Health and Nutrition, Ibaraki, Osaka 567-0085, Japan

**Author notes:** These authors equally contributed to this work (Miyuki Natsume, Mariko Niwa). **Corresponding author**: Yuuki Obata, Ph.D. Laboratory of Intracellular Traffic & Oncology National Cancer Center Research Institute Tsukiji, 5-1-1, Chuo-ku Tokyo, 104-0045, Japan /. **Grant Support** The Japan Society for the Promotion of Science (21K07163 to YO and 22K08883 to TN) Japan Foundation for Applied Enzymology (to YO) Suzuken Memorial Foundation (to YO)<colcnt=2>. **Email** Miyuki Natsume Mariko Niwa Sho Ichikawa Takuma Okamoto Hisazumi Tsutsui Daiki Usukura Takatsugu Murata Ryo Abe Motoyuki Shimonaka Toshirou Nishida Isamu Shiina.

**Keywords:** RTK, KIT, EGFR, MET, GEF, ARF, Golgi/TGN, endoplasmic reticulum, BFA, M-COPA

## Abstract

Normal receptor tyrosine kinases (RTKs) need to reach the plasma membrane (PM) for ligand-induced activation, whereas its cancer-causing mutants can be activated before reaching the PM in organelles, such as the Golgi/*trans*-Golgi network (TGN). Inhibitors of protein export from the endoplasmic reticulum (ER), such as brefeldin A (BFA) and 2-methylcoprophilinamide (M-COPA), can suppress the activation of mutant RTKs in cancer cells, suggesting that RTK mutants cannot initiate signaling in the ER. BFA and M-COPA block the function of ADP-ribosylation factors (ARFs) that play a crucial role in ER–Golgi protein trafficking. However, which ARFs among AFR family proteins are inhibited by BFA or M-COPA, that is, which ARFs are involved in RTKs transport from the ER, remain unclear. In this study, we showed that M-COPA blocked the export of not only KIT but also PDGFRA/EGFR/MET RTKs from the ER. ER-retained RTKs could not fully transduce anti-apoptotic signals, thereby leading to cancer cell apoptosis. Moreover, single knockdown of ARF1, ARF3, ARF4, ARF5, or ARF6 could not block ER export of RTKs, indicating that BFA/M-COPA treatment cannot be mimicked by knockdown of only one ARF member. Interestingly, simultaneous transfection of ARF1, ARF4, and ARF5 siRNAs mirrored the effect of BFA/M-COPA treatment. Consistent with these results, *in vitro* pulldown assays showed that BFA/M-COPA blocked the function of ARF1, ARF4, and ARF5. Taken together, these results suggest that BFA/M-COPA targets at least ARF1, ARF4, and ARF5; in other words, RTKs require the simultaneous activation of ARF1, ARF4, and ARF5 for their ER export.

## Introduction

Ligand-bound receptor tyrosine kinases (RTKs) on the plasma membrane (PM) autophosphorylate tyrosine residues specifically and recruit docking proteins to these phospho-sites to activate downstream signaling cascades, such as the MEK-ERK pathway, the PI3K-AKT pathway, and signal transducer and activator of transcription proteins (STATs), resulting in cell proliferation, survival, and differentiation.^1,2^ Therefore, constitutively active mutants of RTK, such as epidermal growth factor receptor^ΔEX^^19^ (EGFR^ΔEX^^19^), KIT^ΔEX^^11^, and fms-like tyrosine kinase 3 internal tandem duplication (FLT3-ITD), can cause autonomous proliferation of host cells.^3–5^ EGFR, KIT, and FLT3 mutants are well-known as major oncogenic drivers of lung adenocarcinoma (LAD), gastrointestinal stromal tumor (GIST)/mast cell leukemia (MCL), and acute myeloid leukemia (AML), respectively.^3–7^ RTKs are type I transmembrane proteins whose carboxy-terminal tyrosine kinase domain is oriented towards the cytosolic side.^1,2^ Soon after their synthesis in the ER, RTKs undergo partial glycosylation and subsequently move to the Golgi for complex glycosylation, followed by trafficking towards the PM.^8^

Recently, we found that unlike normal RTK, constitutively active RTK mutants, such as KIT and FLT3, are aberrantly retained in the Golgi/*trans*-Golgi network (TGN) and endosome-lysosome compartments, which serve as their oncogenic signaling platforms, leading to autonomous cell growth.^9–13^ In addition, RTKs in the endoplasmic reticulum (ER) are unable to transduce signals,^12–15^ suggesting that blockade of ER export is a new strategy for inhibiting the growth signaling induced by RTK mutants.

ADP-ribosylation factors (ARFs) are small GTPase proteins that serve as a molecular switch of intracellular protein trafficking, such as ER export and endocytosis.^16,17^ ARFs are GTP-bound and active on membranes and GDP-bound and inactive in cytosol.^17^ Guanine nucleotide exchange factors (GEFs) activate ARFs by substituting GDP in ARFs with cytosolic GTP.^16^ Human ARFs and GEFs are composed of 29 and 15 members, respectively.^16,17^ Brefeldin A (BFA) is a well-known inhibitor that blocks protein transport from the ER to the Golgi by immobilizing GEF-ARF complexes.^18–20^ Previously, we reported that KIT and FLT3 cannot move to the Golgi from the ER in the presence of BFA,^9,10,13^ indicating that RTKs moves from the ER in a GEF-ARF-dependent manner. Which GEF-ARF members, however, play a critical role in the biosynthetic transport from the ER remain to be elucidated.

Although the potential of BFA as an anti-cancer drug has been investigated,^21^ its stability is considerably low *in vivo* and its development as a clinical drug has not progressed. On the other hand, 2-methylcoprophilinamide (M-COPA), which similarly blocks the GEF-ARF complex,^22,23^ has higher bioavailability and thus is more efficient *in vivo*.^24^ We recently confirmed using *in vitro* experiments that M-COPA suppresses the growth signaling of KIT/FLT3 mutants in GIST and AML cells by blocking the ER export of these mutants.^12–15^ Therefore, the development of M-COPA and its derivatives as anti-cancer drugs has been initiated currently. Although the mechanism of action is important for establishing these compounds as anti-cancer drugs, which GEF/ARF members are negatively affected by M-COPA are still uncertain. Moreover, inhibition of ER–Golgi transport induces ER stress signaling, which is involved in apoptosis.^25^ It remains unknown whether ER stress signaling is essential for cell growth suppression/apoptosis in cancer cells expressing an RTK mutant, such as KIT in GISTs and EGFR in LAD.

The present study aimed to determine the GEF-ARF members that play a key role in the export of RTKs from the ER through investigating the mechanism of action of M-COPA in inhibiting the growth of RTK-addicted cancer cells. Further, we attempted to identify the GEF and ARF proteins that are affected by M-COPA.

## Results

### M-COPA inhibits the trafficking of RTKs from the ER

To examine whether ER export of RTKs other than KIT was blocked by M-COPA, we used GIST-T1 cells (a human GIST cell line expressing mutant KIT (KIT^Δ560–578^) and wild-type platelet-derived growth factor receptor A (PDGFRA))^26^ and PC-9 cells (a human LAD cell line expressing mutant EGFR (EGFR^Δ746–750^) and wild-type hepatocyte growth factor receptor (MET)). As shown in Figure 1*A*, M-COPA completely suppressed the proliferation of GIST-T1 and PC-9 cells at 1 µM and 200 nM concentrations, respectively. Under control condition, KIT and PDGFRA in GIST-T1 cells were found at the perinuclear region together with Golgi markers (Fig. S1*A*);^10,11,15^ however, these RTKs disappeared from the Golgi region in cells treated with 1 µM M-COPA for 8 h (Fig. 1, *B* and *C*). Golgi matrix protein 130 kDa (GM130) dispersed to the cytosol without any change in its protein levels (Fig. S1, *B* and *C*), indicating that M-COPA affects ER–Golgi trafficking. M-COPA markedly increased the colocalization of KIT/PDGFRA with an ER marker, protein disulfide isomerase (PDI), suggesting that ER export of KIT and PDGFRA is blocked in M-COPA-treated GIST-T1 cells (Fig. 1, *B* and *C*). In PC-9 cells, EGFR and MET were found at the PM, the perinuclear area, and endosomal puncta (Fig. 1, *D* and *E*), as previously reported.^27–30^ Like KIT/PDGFRA in GIST-T1 cells, EGFR and MET were retained in the ER of PC-9 cells in the presence of M-COPA (Fig. 1, *D* and *E*). Immunoblotting showed that these four RTKs were found as doublet bands (Fig. 1, *F* and *G*); the upper bands of KIT, PDGFRA, and EGFR are mature forms, which appear after complex-glycosylation in the Golgi, and MET lower band is a mature form, which is proteolytically cleaved in the TGN.^10,11,31,32^ In M-COPA-treated cells, these RTKs were retained in immature forms unable to be transported from the ER (Fig. 1, *F* and *G*). This trafficking inhibition was correlated with RTK dephosphorylation and decrease of phospho-AKT (pAKT), pERK, and pSTATs, indicating that these RTKs could not be completely activated in the ER. As shown in Figure S1*D*, an immaturely glycosylated TGN46 protein was also increased in M-COPA-treated cells, suggesting that M-COPA blocks the general protein export from the ER.

**Figure 1.**
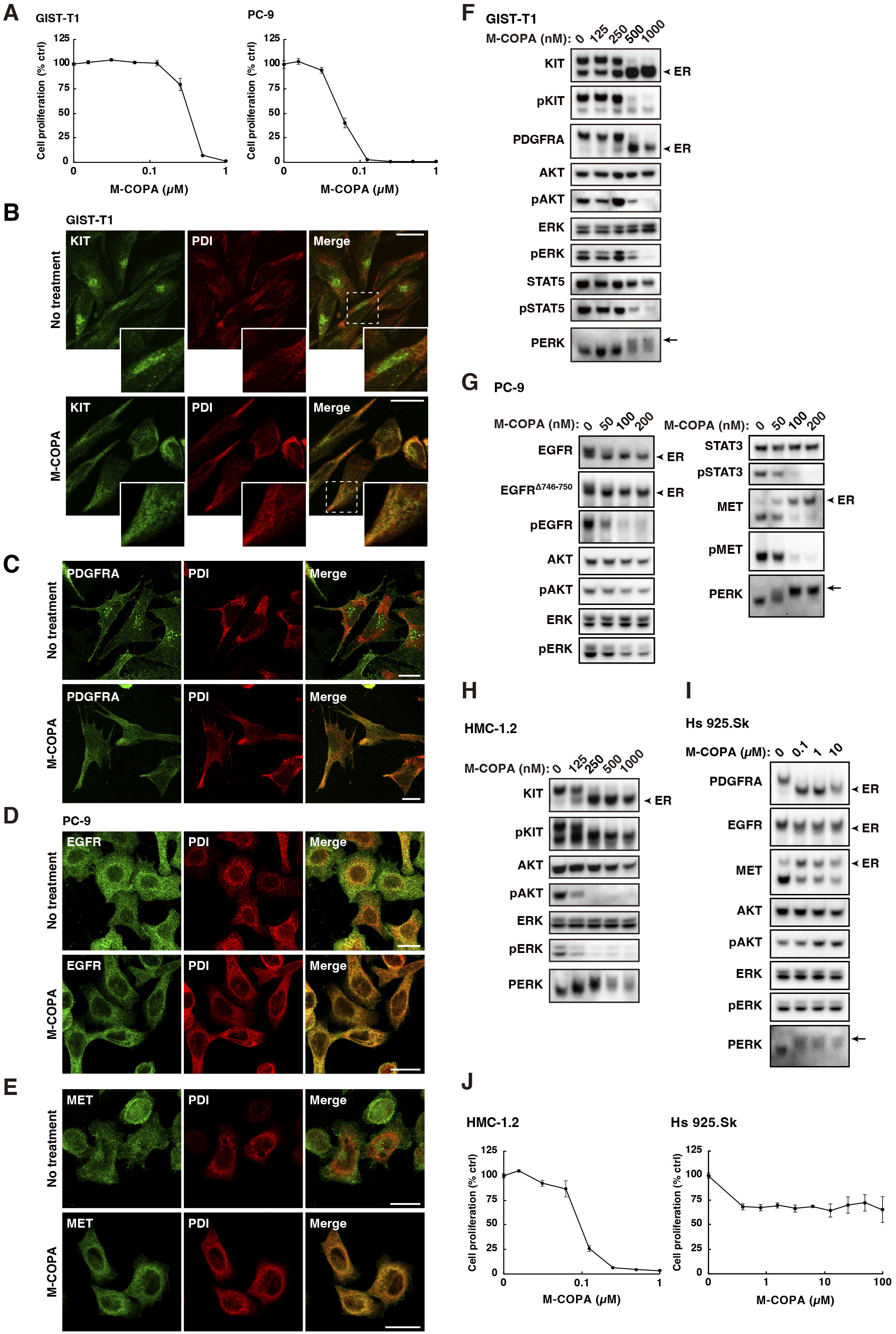
M-COPA inhibits ER export of RTKs, such as KIT, PDGFRA, EGFR, and MET. *A,* GIST-T1 and PC-9 cells were treated with M-COPA for 48 h. Cell proliferation was assessed by ATP production. Values represent mean ± SD (*n* = 3). *B–I,* GIST-T1 (*B*, *C*, and *F*), PC-9 (*D*, *E*, and *G*), HMC-1.2 (*H*), and Hs 925.Sk cells (*I*) were treated with M-COPA for 8 h. *B–E*, Cells were immunostained for protein disulfide isomerase (PDI, ER marker) in conjunction with KIT, PDGFRA, EGFR, or MET. Magnified images of the boxed area are shown. Bars, 20 µm. *F–I,* Lysates were immunoblotted with the indicated antibody. Arrowheads and arrows indicate ER-retained RTKs and phosphorylated PERK, respectively. Note that RTKs were retained in the ER in M-COPA-treated cells. *J,* HMC-1.2 and Hs 925.Sk cells were treated with M-COPA for 48 h. Cell proliferation was assessed by ATP production. Values represent mean ± SD (*n* = 3).

The blockade of ER–Golgi trafficking induces ER stress responses, such as PKR-like ER kinase (PERK) activation.^25^ PERK was converted to its higher molecular weight form because of multiple phosphorylations in the presence of M-COPA (Fig. 1, *F* and *G*), indicating that M-COPA affects the secretory pathway from the ER. Similar results were observed in a MCL cell line, HMC-1.2, which expresses an active KIT mutant (Fig. 1*H*). In addition to cancer cells, a human skin fibroblast cell line Hs 925.Sk cells also contained immature RTKs in the presence of M-COPA (Fig. 1*I*). Interestingly, pAKT and pERK in the normal cells were unaffected. Indeed, compared with that of cancer cells, the growth sensitivity of Hs 925.Sk to M-COPA was quite low (Fig. 1*J*), indicating that Hs 925.Sk cells can activate growth signaling without PM-localized RTKs. Taken together, these results suggest that M-COPA inhibits the ER export of not only KIT but also PDGFRA, EGFR, and MET.

### M-COPA can induce apoptosis in a PERK-independent manner in RTK-mutated cells

Caspase-3 was cleaved in GIST-T1 and HMC-1.2 cells treated with M-COPA for 24 h and 8 h, respectively, which is a sign of apoptosis (Fig. 2*A*). Previous studies have demonstrated that ER stress-induced PERK activation leads host cells to apoptosis.^25^ Thus, we investigated whether the induction of PERK activation is necessary for M-COPA-induced apoptosis using PERK inhibitor II (PERKi II). As shown in Figure 2*B* and 2*C*, PERKi II completely suppressed the M-COPA-induced mobility shift of PERK bands, confirming that it blocks PERK activation. Interestingly, under these conditions, caspase-3 was also cleaved, suggesting that growth signaling inhibition through the blockade of ER export of mutant KIT is sufficient for M-COPA-induced apoptosis of GIST-T1 and HMC-1.2 cells. Moreover, immunoblotting results revealed that the protein levels of AKT and STAT5 were decreased, and fragments of these proteins were present in the lower molecular weight region (Fig. 2*D*). We suspected that these proteins were cleaved by caspases. When cells were treated with M-COPA plus a pan-caspase inhibitor, Z-VAD-FMK,^33^ AKT and STAT5 cleavage were suppressed (Fig. 2*D*). The effect of Z-VAD-FMK was confirmed by the reduction of completely cleaved caspase-3. Z-VAD-FMK addition could not restore the protein ERK levels decreased by M-COPA, indicating that caspases that are activated by M-COPA specifically cleave AKT and STAT5. Taken together, these results suggest that caspase-3 activation by M-COPA is independent of PERK activation but dependent on the blockade of ER export of mutant RTKs in cancer cells.

**Figure 2.**
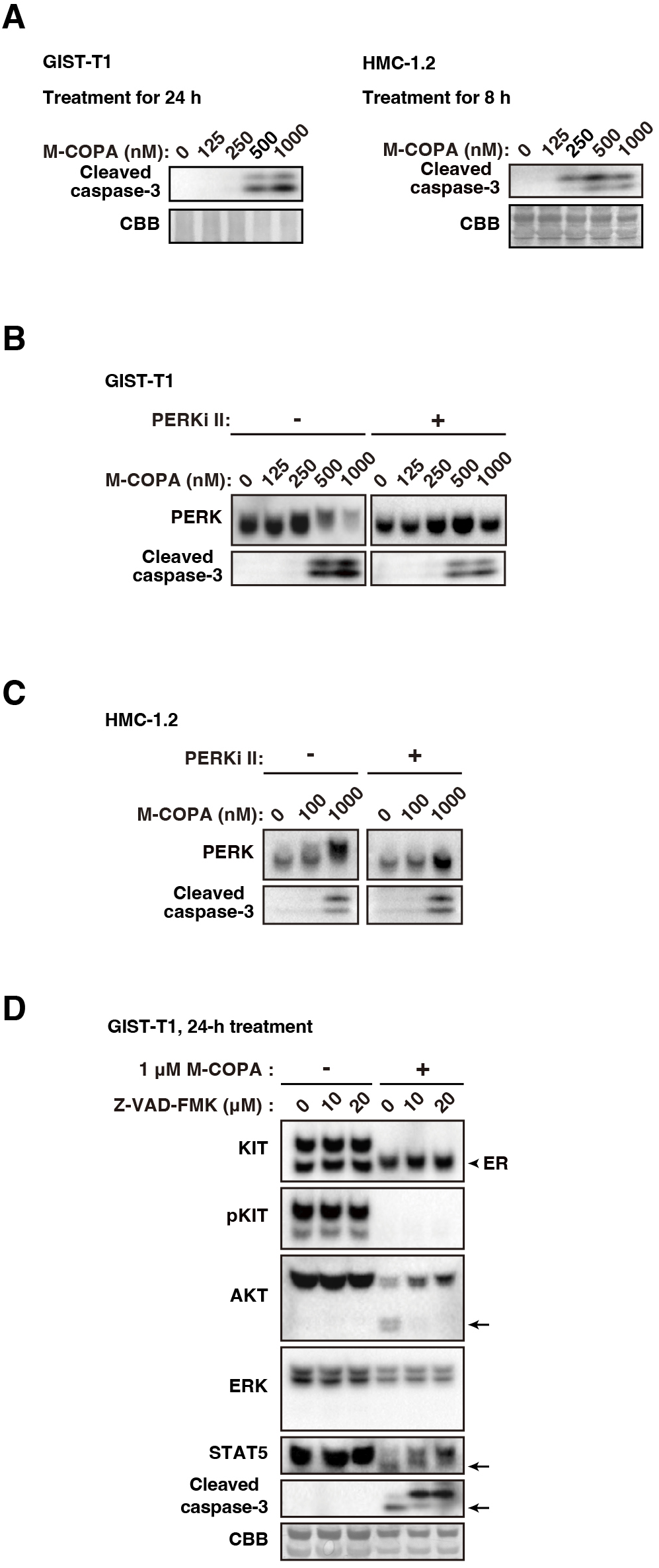
M-COPA induces apoptosis PERK-independent manner in GIST-T1 and HMC-1.2 cells. *A,* GIST-T1 (left) and HMC-1.2 cells (right) were treated with M-COPA for 24 h and then immunoblotted for cleaved caspase-3. Total protein levels were confirmed by Coomassie Brilliant Blue (CBB) staining. *B,* GIST-T1 cells were treated with M-COPA and/or PERK inhibitor II (PERKi II) for 24 h and then immunoblotted for PERK and cleaved caspase-3. *C,* HMC-1.2 cells were treated with M-COPA and/or PERKi II for 8 h and immunoblotted. *D,* GIST-T1 cells were treated with M-COPA and/or Z-VAD-FMK (pan-caspase inhibitor) for 24 h and immunoblotted. An arrowhead indicates ER-retained KIT. Arrows indicate the cleaved AKT, STAT5, and caspase-3. Total protein levels were confirmed by CBB staining.

### Simultaneous inhibition of ARF1, ARF4, and ARF5 is required for mimicking the action of BFA/M-COPA

Several studies have reported that BFA, golgicide A, and M-COPA block intracellular trafficking by suppressing the dissociation of ARF from GEF.^18–20,34,35^ Therefore, we explored whether knockdown of GEF and ARF can phenocopy BFA/M-COPA treatment to understand the precise targets of these inhibitors. Golgi-specific BFA-resistance guanine nucleotide exchange factor 1 (GBF1), BFA-inhibited guanine nucleotide-exchange protein 1 (BIG1), and BIG2 are widely known as BFA-sensitive GEFs.^36^ However, knockdown of GBF1, BIG1, or BIG2 did not exhibit similar results as shown by BFA/M-COPA treatment in the mobility shift of KIT/PDGFRA bands, signal inhibition, or PERK activation (Fig. 3*A*). Although simultaneous knockdown of GBF1, BIG1, and BIG2 slightly decreased the KIT level, pERK, and pSTAT5, their knockdown did not yield similar effects to that of BFA/M-COPA treatment (Fig. 3*B*), suggesting that the action of BFA/M-COPA cannot be explained by GBF1/BIG1/BIG2 inhibition alone. In other words, in addition to GBF1/BIG1/BIG2, other GEFs play a role in RTK trafficking from the ER. At present, identification of BFA/M-COPA-sensitive GEFs is under way.

**Figure 3.**
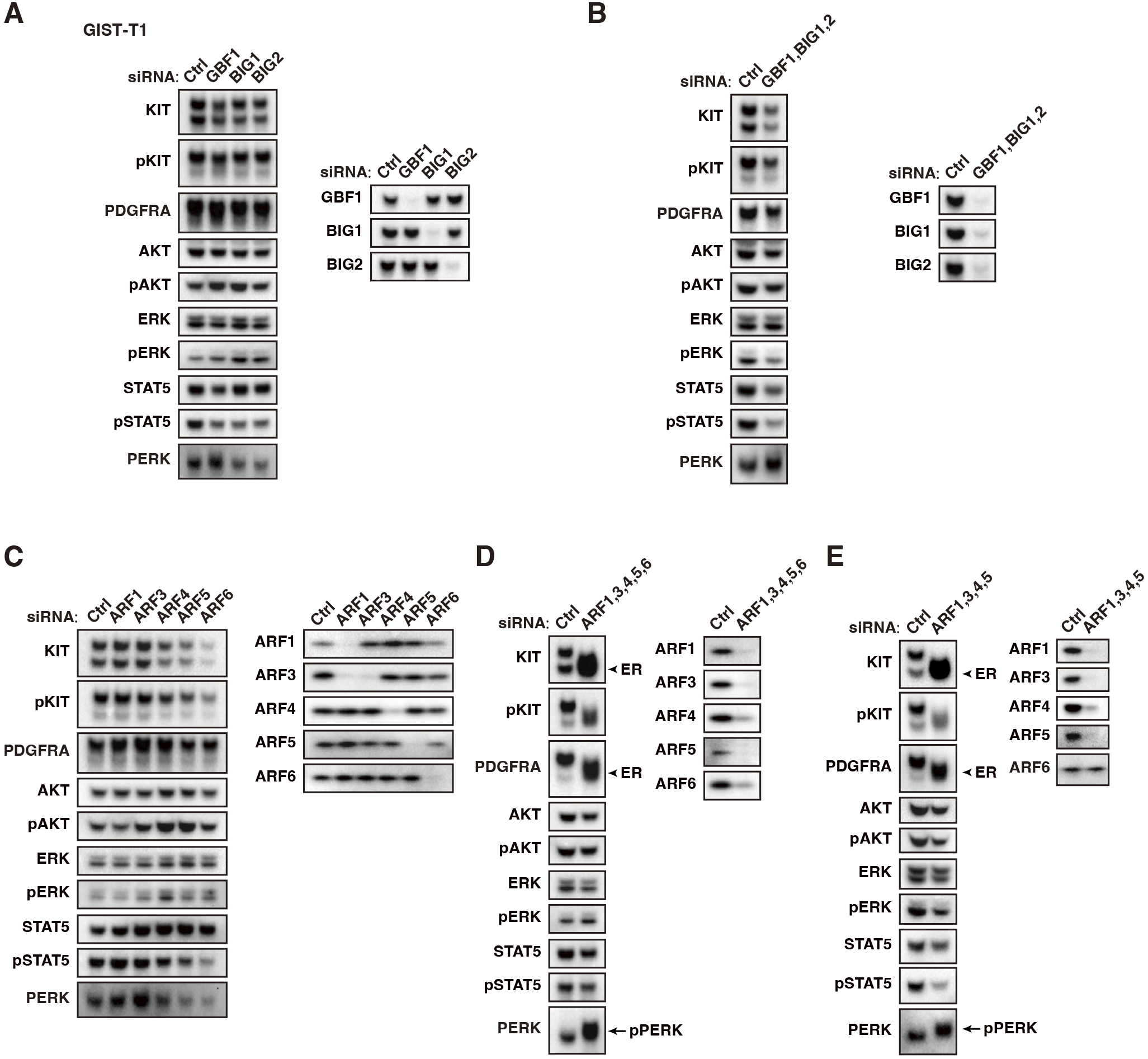
Simultaneous knockdown of ARFs is required for mimicking BFA/M-COPA treatment in GIST-T1 cells. *A-E,* GIST-T1 cells were transfected with the indicated siRNA for 48 h. Lysates were immunoblotted with the indicated antibodies. Arrowheads and arrows indicate ER-retained RTKs and phosphorylated PERK (pPERK), respectively.

Next, we knocked down major ARF members, such as ARF1, ARF3, ARF4, ARF5, and ARF6. Like the results observed after GEF knockdown, single knockdown of an ARF member did not phenocopy M-COPA treatment (Fig. 3*C*). Although knockdown of ARF4, ARF5, and ARF6 decreased the levels of KIT and pSTAT5, the effect was not due to inhibition of ER export, as the accumulation of immature RTKs was not observed. Notably, ARF1 siRNA greatly reduced the ARF3 protein levels. We tested other ARF1 siRNAs, which also reduced ARF3 (Fig. S2*A*). Thus, we concluded that ARF3 reduction is not due to the off-target effect of ARF1 siRNA. As shown in Figure S2*B*, M-COPA decreased the ARF3 level, indicating that ARF1 may play a role in the stability of ARF3 protein. Next, we simultaneously knocked down ARF1, ARF3, ARF4, ARF5, and ARF6. Interestingly, simultaneous knockdown of these ARFs showed the band shift of KIT, PDGFRA, and PERK activation, indicating that these RTKs are retained in the ER by ARF1–6 knockdown (Fig. 3*D*). ARF6 siRNA was not required for the blockade of ER export of the RTKs (Fig. 3*E*), suggesting that ARF6 is not involved in ER–Golgi trafficking.

We further determined the siRNA required for mimicking BFA/M-COPA treatment. As shown in Figure 4*A*, ER export of RTKs was not inhibited in the absence of ARF1 or ARF4 siRNA. ARF5 siRNA addition to ARF1/ARF4 siRNAs expedited the effect of BFA/M-COPA treatment (Fig. 4, *A* and *B*). We could not conclude whether ARF3 knockdown was required for phenocopying BFA/M-COPA treatment, because ARF1 knockdown decreased ARF3 levels, as shown in Figures 3*C* and S3*A*. Unlike M-COPA treatment, the simultaneous knockdown did not affect pAKT or pERK. A previous study showed that ARF inhibition with a dominant negative ARF mutant induces feed forward stimulation of GBF1.^37^ We expected that upon ARF1/4/5 knockdown, GBF1 would be recruited to the membrane, causing ARFs other than ARF1/4/5 to be activated as compensation. Therefore, in addition to ARF1/4/5 siRNAs, we transfected GBF1 siRNA to mimic BFA/M-COPA treatment. As shown in Figure 4*C* and 4*D*, most effects of ARF1/4/5 siRNAs plus GBF1 siRNA were similar to that of BFA/M-COPA treatment: KIT and PDGFRA were retained in an immature form in the ER, KIT signaling was inhibited, and PERK was activated. However, unlike M-COPA treatment, the knockdown did not induce dispersion of GM130 (Fig. 4*D*, lower panels; compare with Fig. S1*B*), indicating that additional knockdown of ARFs/GEFs may be required for completely mimicking the BFA/M-COPA treatment. As shown in Figure 4*E*, ARF1/4/5 and GBF1 knockdown in PC-9 cells demonstrated similar results in that ER export of RTKs was blocked, resulting in inactivation of EGFR, MET, AKT, ERK, and STAT3. As shown in Figure S2*C*, the simultaneous knockdown increased immature TGN46 protein, suggesting that ARF1/4/5 and GBF1 plays an essential role not only in the trafficking of RTK but also in that of other membrane-bound proteins. Taken together, these results suggest that the simultaneous inhibition of at least ARF1, ARF4, ARF5, and GBF1 is required for mimicking the effect of BFA/M-COPA treatment; in other words, RTKs require the simultaneous activation of ARF1, ARF4, ARF5, and GBF1 for their ER export.

**Figure 4.**
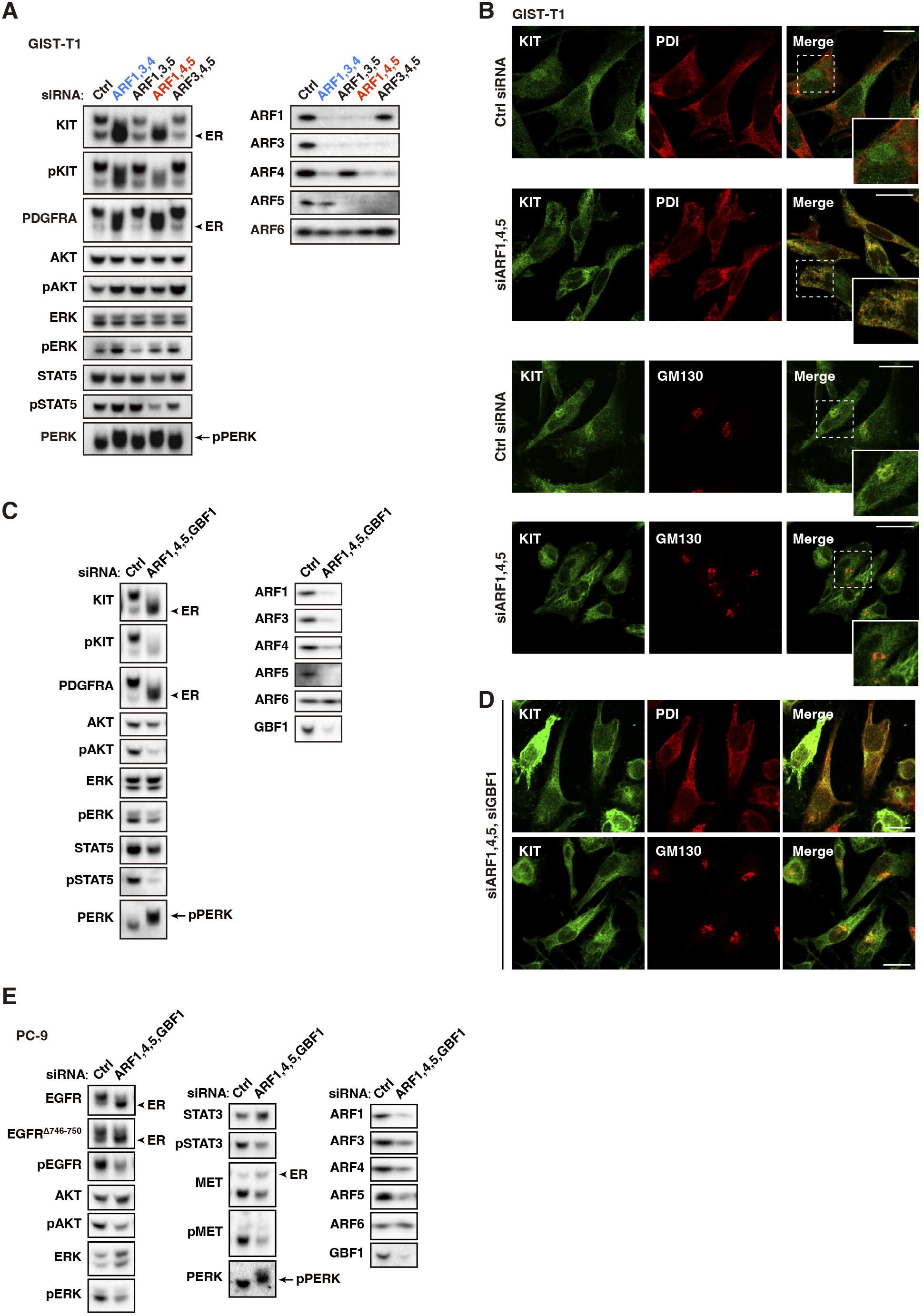
At least simultaneous knockdown of ARF1, ARF4, and ARF5 is necessary for mimicking BFA/M-COPA treatment. *A–D,* GIST-T1 cells were transfected with the indicated siRNAs and then cultured for 48 h. *A* and *C,* Lysates were immunoblotted with the indicated antibodies. *B* and *D,* Cells were immunostained for KIT, PDI (ER marker), and Golgi matrix protein 130 kDa (GM130). Magnified images of the boxed area are shown. Bars, 20 µm. Note that transfection of ARF1, ARF4, ARF5, and GBF1 siRNAs got closer to the effect of BFA/M-COPA treatment. *E,* PC-9 cells were transfected with the indicated siRNAs for 24 h and then immunoblotted. Arrowheads and arrows indicate ER-retained RTKs and phosphorylated PERK (pPERK), respectively.

### M-COPA inhibits activation and membrane localization of ARF1, ARF4, and ARF5

Next, we investigated whether M-COPA inhibits ARF1/4/5. We performed a pulldown assay with glutathione S-transferase-fused Golgi-localized γ-adaptin ear-containing, ARF-binding protein 3 (GST-GGA3), which binds to active ARF.^38,39^ As shown in Figure 5*A*, ARF1, ARF4, ARF5, and ARF6 were activated in GIST-T1 cells, although ARF3 pulldown was not detected probably because of a characteristic of ARF3 antibody. Interestingly, M-COPA decreased the levels of GGA3-bound ARF1, ARF4, and ARF5. ARF6 binding was not affected, suggesting that M-COPA specifically inhibits the activation of ARF1, ARF4, and ARF5. These pulldown assay data support the results of knockdown that ARF6 is not involved in ER export of RTKs. BFA treatment results were consistent with those of M-COPA treatment (Fig. 5*B*), suggesting that both these compounds can inhibit ARF1, ARF4, and ARF5 but not ARF6. Furthermore, M-COPA also inhibited ARF1, ARF4, and ARF5 in PC-9 cells (Fig. 5*C*).

**Figure 5.**
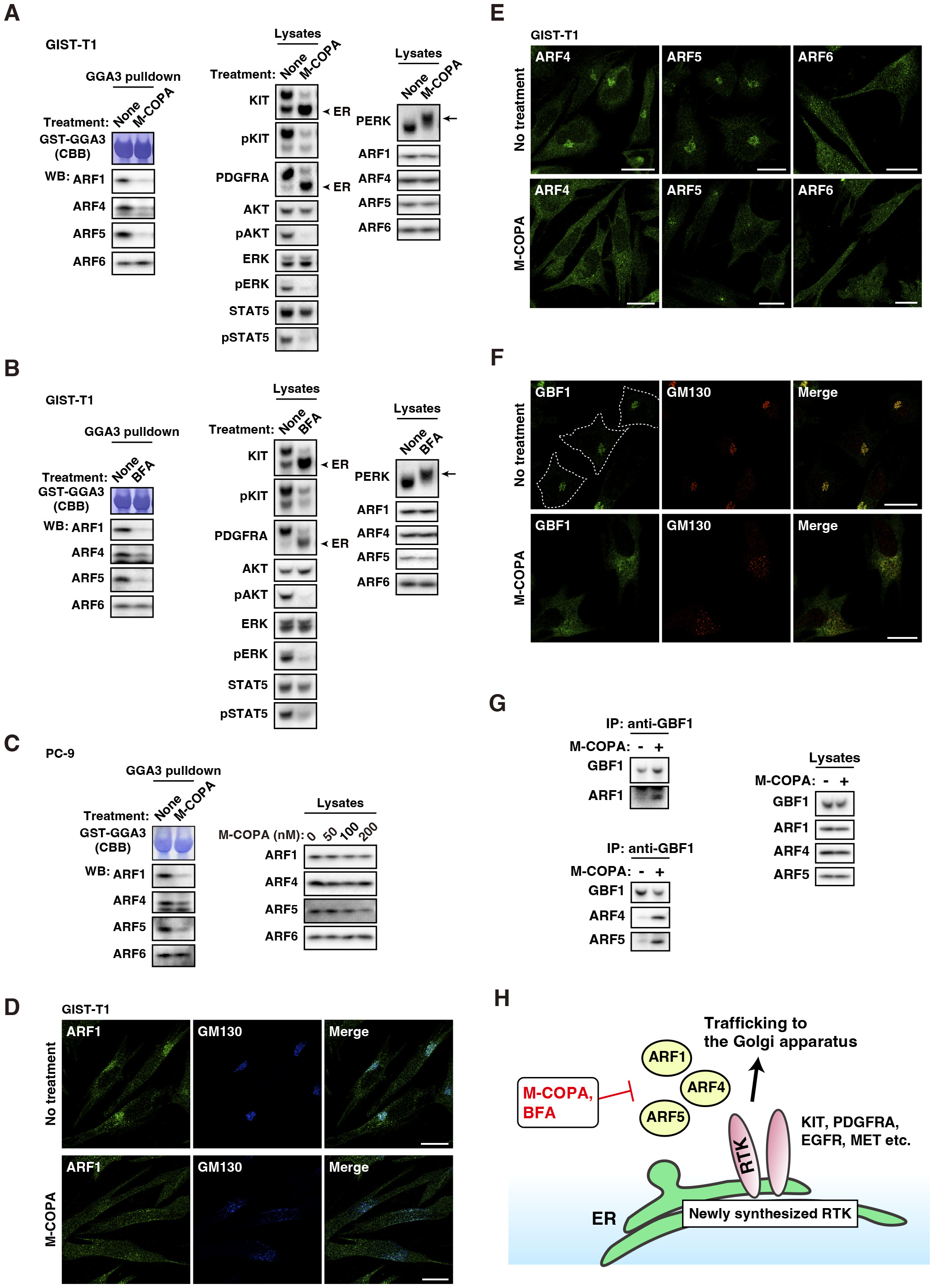
M-COPA and BFA inhibit the activation of ARF1, ARF4, and ARF5 but not ARF6. *A* and *B,* GIST-T1 cells were treated with (*A*) 1 µM M-COPA or (*B*) 1 µM BFA for 4 h. ARFs were pulled down with GST-tagged GGA3(1-316) (GST-GGA3) and then immunoblotted (left). Amounts of GST-GGA3 were confirmed by CBB staining. Lysate immunoblots are shown in the center and the right side. Arrowheads and arrows indicate ER-retained RTKs and phosphorylated PERK, respectively. *C,* PC-9 cells were treated with 200 nM M-COPA for 8 h. ARFs were pulled down with GST-GGA3 and then immunoblotted. Lysate immunoblots under the same condition are shown in the right side and Figure 1G. *D–F,* GIST-T1 cells treated with 1 µM M-COPA for 4 h were immunostained with the indicated antibody. Dashed lines indicate cell borders. Bars, 20 µm. PDI, ER marker; GM130, Golgi marker. *G,* GIST-T1 cells were treated with 1 µM M-COPA for 4 h. GBF1 was immunoprecipitated and then immunoblotted. IP, immunoprecipitation. *H,* A schematic model of the ARF-dependent trafficking of RTKs from the ER. BFA and M-COPA inhibit the dissociation of ARF1, ARF4, ARF5 from GBF1, resulting in blockade of ER export of RTKs, such as KIT, PDGFRA, EGFR, and MET. Therefore, RTKs are exported from the ER in an ARF1/4/5-dependent manner. In addition, RTK mutants cannot become fully active form, thus BFA and M-COPA inhibit the growth signaling in RTK-addicted cancer cells.

Next, we confirmed the localization of ARFs in M-COPA-treated cells. Previous studies have reported that active ARFs are recruited to organelles, whereas inactive ARFs are dissociated from the membranes to the cytosol.^16,40,41^ Under normal condition, ARF1, ARF4, and ARF5 were mainly found in the perinuclear region (Figs. 5, *D* and *E* and S3). However, they relocated to the cytosol after M-COPA treatment, but ARF6 distribution was not affected by the treatment. These results supported the data of the GST pulldown assay. Next, we verified whether GBF1 activity was decreased in M-COPA-treated cells. Similar to ARF1/4/5, perinuclear GBF1 disappeared after M-COPA treatment (Fig. 5*F*). Because BFA immobilizes the ARF-GBF1 complexes,^20,34,35^ we performed co-immunoprecipitation assay to examine whether M-COPA affected the association of GBF1 with ARFs. As shown in Figure 5*G*, co-immunoprecipitation of ARF1/ARF4/ARF5 with GBF1 was greatly increased by M-COPA treatment, indicating that GBF1 was indeed inhibited by the treatment. Taken together, these results suggest that BFA/M-COPA simultaneously inhibits ARF1, ARF4, and ARF5 by immobilizing their association with GBF1.

## Discussion

In this study, we demonstrated that simultaneous inhibition of at least ARF1, ARF4, and ARF5 is required for mimicking BFA/M-COPA treatment. In the knockdown condition in GIST cells, KIT mutant and PDGFRA are retained in the ER, where they are unable to reach their fully active form. EGFR mutant and MET in LAD cells are exported from the ER in a similar manner to KIT. These results suggest that these RTKs are exported from the ER towards the Golgi in a ARF1/4/5-dependent manner (Fig. 5*H*). Because knockdown of only one ARF member does not mimic the effect of BFA/M-COPA, simultaneous inhibition of ARF1, ARF4, and ARF5 is essential for investigating the mechanism of action of these drugs.

Recent studies have reported that RTKs other than KIT, such as FLT3-ITD, PDGFRA, MET, and IGF-1R, also use the Golgi/TGN as their signaling platform in cancer cells.^13,30,42–44^ Moreover, non-RTK proto-oncogene products, such as Src-family kinases, RAS, and mTOR, can initiate growth signaling from the Golgi/TGN.^10,45–49^ Therefore, these studies raise a possibility that inhibitors of protein trafficking from the ER to the Golgi can be potential candidates for anti-cancer drugs. Although the bioavailability of M-COPA is higher than that of BFA, the stability of M-COPA *in vivo* is not sufficient for a clinical trial. Thus, development of M-COPA derivatives with considerably improved *in vivo* stability is now under way.

BFA and M-COPA cause PERK activation in GIST-T1 and HMC-1.2 cells, but the activation is not necessary for inducing apoptosis. In these cells, mutant KIT-dependent AKT activation has an essential role in the anti-apoptotic status;^14,50^ thus, AKT inhibition through blockade of ER export of KIT is sufficient for inducing apoptosis. In addition to PERK, ATF6 and IRE1 pathways are known to be activated by ER stress.^25^ However, these ER stress-induced activations may not be essential for RTK-addicted cancer cells.

M-COPA treatment for a long period induced caspase-dependent limited proteolysis of AKT and STAT5 in GIST-T1 cells. The cleavage causes these proteins to lose survival signaling.^51^ Previous studies have shown that over 1,000 proteins including AKT are cleaved by caspases.^52,53^ In our study, cleaved STAT5 was present in M-COPA-treated cells, indicating that STAT5 is also a substrate candidate for caspases. Because AKT activation is essential for anti-apoptosis signaling in mutant KIT-addicted cancer cells,^14,50^ the cleavage of AKT may serve as a positive-feedback mechanism for M-COPA-induced apoptosis. Thus, once caspases are activated, intracellular AKT phosphorylation levels may be synergistically decreased in M-COPA-treated cells.

Our study showed that the growth sensitivity of normal cells to M-COPA is markedly lower than that of RTK-addicted cancer cells. M-COPA blocked ER export of EGFR, MET, and PDGFRA in Hs 925.Sk cells but could not inhibit AKT and ERK activation. We previously observed ligand-dependent KIT signaling in a normal mast cell line treated with M-COPA.^15^ Considering that M-COPA derivatives are being developed as an anti-cancer drug, the difference of sensitivity to M-COPA between cancer cells and normal cells is useful to avoid side effects.

Molecular targeted drugs, such as a tyrosine kinase inhibitor (TKI) imatinib, can extend the life of patients with progressive GIST.^54,55^ However, patients can develop resistance to imatinib within 2–3 years after treatment. Secondary mutations in the *KIT* gene, which causes KIT to lose sensitivity to TKIs, are frequently found in imatinib-resistant GISTs. TKI resistance is found in EGFR-mutated LAD in a similar manner to KIT.^3,56^ In this study, M-COPA inhibits KIT signaling of HMC-1.2 cells endogenously expressing KIT^D816V^, which is an imatinib-resistant mutant. Therefore, trafficking inhibition from the ER would be a promising strategy for the suppression of growth signaling through TKI-resistant RTK mutants.

Human ARF3 is known as a BFA-sensitive ARF member, which is structurally similar to ARF1; both are classified as class I ARFs.^16,36^ However, we could not conclude in this study whether ARF3 knockdown is essential for mimicking BFA/M-COPA treatment, because ARF1 depletion decreases the protein level of ARF3. Moreover, ARF3 antibodies used in this study are ineffective in GST-GGA3 pulldown assays and immunofluorescence. Further studies are required for elucidating the role of ARF3 in protein export from the ER.

Previous studies as well as this present study showed that BFA and M-COPA can disperse GM130 from the Golgi, indicating that these compounds affect the *cis*-side of the Golgi apparatus.^35,57^ Although simultaneous knockdown of ARF1, ARF4, ARF5, and GBF1 blocks protein trafficking from the ER, the knockdown does not disperse GM130 from the Golgi area. A previous study reported that ARF4 but not ARF1/ARF5 plays a role in BFA-induced growth inhibition,^58^ indicating that ARF4 protein also influences the effects of BFA after treatment. This may be a reason why the simultaneous knockdown did not completely mimic BFA/M-COPA treatment. Furthermore, simultaneous knockdown of GBF1, BIG1, and BIG2 also did not mimic BFA/M-COPA treatment. Knockdown of additional GEFs may be required for understanding the mode of action of BFA/M-COPA. Thus, analyses of “additional hits” of GEFs and ARFs for fully understanding BFA/M-COPA action is currently in motion.

In conclusion, simultaneous inactivation of at least ARF1, ARF4, and ARF5 is required for mimicking BFA/M-COPA treatment. Our results provide evidence that M-COPA certainly inhibits ARF1, ARF4, and ARF5 but not ARF6. Moreover, from a clinical perspective, our study can contribute to the development of M-COPA and its derivatives as a molecular targeted drug that suppresses the growth of RTK-addicted cancers. Therefore, our study provides significant insights into intracellular trafficking in cancer cell biology and for molecular targeted therapeutics.

## Experimental procedures

### Cell culture

GIST-T1 (Cosmo Bio, Tokyo, Japan)^26^ and Hs 925.Sk cells (American Type Culture Collection, Manassas, VA, USA) were cultured at 37°C in Dulbecco’s Modified Eagle’s Medium (DMEM) supplemented with 10% fetal calf serum (FCS), penicillin, and streptomycin (Pen/Strep). PC-9 (The European Collection of Authenticated Cell Cultures, Salisbury, UK) and HMC-1.2 cells^59^ were cultured at 37°C in RPMI1640 supplemented with 10% FCS, and Pen/Strep. To culture HMC-1.2 cells, 50 µM 2-mercaptoethanol was added. All human cell lines were tested for *Mycoplasma* contamination using a MycoAlert Mycoplasma Detection Kit (Lonza, Basel, Switzerland).

### Cell proliferation assay

Cells were cultured with M-COPA for 48 h. Cell proliferation was quantified using the CellTiter-Glo Luminescent Cell Viability Assay (Promega, Madison, WI, USA), according to the manufacturer’s instructions. ATP production was measured with the 2030 ARVO X3 Multilabel Plate Reader (PerkinElmer) or Synergy H1 Multimode Microplate Reader (Agilent, Santa Clara, CA, USA).

### Antibodies

The lists of antibodies used for immunoblotting, immunoprecipitation, and immunofluorescence are shown in Supplementary Tables 1–3.

### Chemicals

Z-VAD (Ome)-FMK (Abcam, Cambridge, UK), PERK inhibitor II (Sigma-Aldrich, St. Louis, MO, USA), imatinib mesylate (Cayman Chemical, Ann Arbor, MI, USA), and M-COPA^22,23^ were dissolved in DMSO. BFA (Sigma-Aldrich) was dissolved in ethanol.

### Gene silencing with siRNA

To silence *ARF* and *GEF* genes, ON-TARGETplus SMARTpool siRNAs and Silence Select Pre-Designed siRNAs were purchased from Horizon Discovery (Waterbeach, UK) and Thermo Fisher Scientific (Rockford, IL, USA), respectively. A list of the siRNAs used is provided in Supplementary Table 4. Electroporation was performed using the NEON Transfection System (Thermo Fisher Scientific), according to the manufacturer’s instructions.

### Immunofluorescence confocal microscopy

GIST-T1 or PC-9 cells were cultured on poly L-lysine-coated coverslips and fixed with 4% paraformaldehyde for 20 min at room temperature. The fixed cells were permeabilized and blocked for 30 min in Dulbecco’s phosphate-buffered saline (D-PBS(-)) supplemented with 0.1% saponin and 3% bovine serum albumin (BSA) and then incubated with primary and secondary antibodies for 1 h each. After washing with D-PBS(-), the cells were mounted with Fluoromount (Diagnostic BioSystems, Pleasanton, CA, USA). Confocal images were obtained with a FLUOVIEW FV10i (Olympus, Tokyo, Japan) or TCS SP5 II/SP8 (Leica, Wetzlar, Germany) laser scanning microscope. Composite figures were prepared using FLUOVIEW FV1000 Viewer (Olympus), Leica Application Suite X Software (Leica), Photoshop, and Illustrator software (Adobe, San Jose, CA, USA).

### Western blotting

Lysates prepared in sodium dodecyl-sulfate polyacrylamide gel electrophoresis (SDS-PAGE) sample buffer were subjected to SDS-PAGE and electrotransferred onto polyvinylidene fluoride membranes. Briefly, 5% skim milk in Tris-buffered saline with Tween 20 (TBST) was used to dilute the antibodies. For immunoblotting with anti-pKIT, anti-pEGFR, or anti-pMET, the antibodies were diluted with 3% BSA in TBST. Immunodetection was performed using Immobilon Western Chemiluminescent HRP Substrate (Sigma-Aldrich). Sequential re-probing of membranes was performed after complete removal of antibodies with Restore PLUS Western Blot Stripping Buffer (Thermo Fisher Scientific) or inactivation of peroxidase by 0.1% NaN_3_. The results were analyzed using ChemiDoc XRC+ with Image Lab software (Bio-Rad, Hercules, CA, USA). To immunoblot for cleaved caspase-3 of GIST-T1 cells, cells that are detached from the bottom of dish and adherent cells were collected, then lysed. Total protein levels were confirmed by Coomassie Brilliant Blue (CBB) staining.

### Immunoprecipitation

Lysates from 2–4 x 10^6^ cells were prepared in NP-40 lysis buffer (50 mM HEPES pH 7.4, 10% glycerol, 0.1–1% NP-40, 4 mM EDTA, 100 mM NaF, 1 mM Na_3_VO_4_, protease inhibitor cocktail, 2 mM β-glycerophosphate, 2 mM sodium pyrophosphate, and 1 mM phenylmethylsulfonyl fluoride). Immunoprecipitation was performed at 4°C for 5 h using protein G Dynabeads precoated with anti-GBF1 antibody. The immunoprecipitates were dissolved in SDS-PAGE sample buffer then immunoblotted.

### GST-GGA3(1–316) pulldown assay

Lysates from 2–4 × 10^6^ cells were prepared in lysis buffer supplemented with 10 mM NaF, 1 mM Na_3_VO_4_, protease inhibitor cocktail, 2 mM β-glycerophosphate, 2 mM sodium pyrophosphate, and 1 mM phenylmethylsulfonyl fluoride. According to the manufacturer’s instructions (Cytoskeleton, Denver, CO, USA), ARFs in cell lysates were incubated with GST-GGA3(1–316)-coated Sepharose beads for 1 h at 4L. These beads were washed once, then added SDS-PAGE sample buffer. Eluted proteins were resolved by SDS-PAGE and immunoblotted. Amounts of GST-GGA3(1–316) were checked by CBB staining.

## Acknowledgments

This work was supported by a grant-in-aid for Scientific Research from the Japan Society for the Promotion of Science, Japan (21K07163 to Y. O. and 22K08883 to T. N.), a research grant from the Japan Foundation for Applied Enzymology, Japan (to Y. O.), and the Suzuken Memorial Foundation, Japan (to Y. O.).

## Author contributions

Miyuki Natsume, Mariko Niwa, S. I., and T. O. investigation; Miyuki Natsume, Mariko Niwa, M. S., and T. N. data interpretation; I. S., T. N., and Y. O. conceptualization; I. S., M. S., T. N., and Y. O. supervision; T. N., and Y. O. funding acquisition; I. S., and Y. O. project administration; T. M., H. T., D. U., M. S., and I. S. resources; Miyuki Natsume, Mariko Niwa, and Y.O. writing-original draft; Miyuki Natsume, Mariko Niwa, M. S., R.A., T. N., I. S., and Y. O. writing–review and editing

## Conflicts of interest

The authors declare that they have no conflicts of interest with the contents of this article.

### Abbreviations

AML: acute myelogenous leukemia
ARF: ADP-ribosylation factor;
BFA: brefeldin A; BIG1/2, BFA-inhibited guanine nucleotide-exchange protein 1/2
CBB: Coomassie Brilliant Blue
ER: endoplasmic reticulum
ex: exon
ERK: extracellular signal-regulated kinase
FLT3-ITD: FMS-like tyrosine kinase 3-intenal tandem duplication
GBF1: Golgi-specific BFA-resistance guanine nucleotide exchange factor 1
GEF: guanine nucleotide exchange factor
GGA: Golgi-associated, γ-adaptin ear-containing, ARF-binding protein
GIST: gastrointestinal stromal tumor
GM130: Golgi matrix protein 130 kDa
IP/IPs: immunoprecipitation/immunoprecipitates
LAD: lung adenocarcinoma
MCL: mast cell leukemia
M-COPA: 2-methylcoprophilinamide
PDGFR: platelet-derived growth factor receptor
PDI: protein disulfide isomerase
PERK: protein kinase R-like endoplasmic reticulum kinase
pKIT: phospho-KIT
PM: plasma membrane
RTK: receptor tyrosine kinase
siRNA: small interfering RNA
STAT: signal transducer and activator of transcription
TGN: *trans-*Golgi network
TKI: tyrosine kinase inhibitor

## References

1. Lemmon, M. A., and Schlessinger, J. (2010). Cell signaling by receptor tyrosine kinases. Cell 141, 1117–1134

2. Du, Z., and Lovly, C. M. (2018). Mechanisms of receptor tyrosine kinase activation in cancer. Mol. Cancer 17, 58

3. Linardou, H., Dahabreh, I. J., Bafaloukos, D., Kosmidis, P., and Murray, S. (2009). Somatic EGFR mutations and efficacy of tyrosine kinase inhibitors in NSCLC. Nat. Rev. Clin. Oncol. 6, 352–366

4. Lennartsson, R., and Rönnstrand, L. (2012). Stem cell factor receptor/c-Kit: from basic science to clinical implications. Physiol. Rev. 92, 1619–1649

5. Kazi, J. U., and Rönnstrand, L. (2019). FMS-like Tyrosine Kinase 3/FLT3: From Basic Science to Clinical Implications. Physiol. Rev. 99, 1433–1466

6. Hirota, S., Isozaki, K., Moriyama, Y., Hashimoto, K., Nishida, T., Ishiguro, S., et al. (1998). Gain-of-function mutations of *c-kit* in human gastrointestinal stromal tumors. Science 279, 577–580

7. Boissan, M., Feger, F., Guillosson, J. J., and Arock, M. (2000). c-Kit and c-kit mutations in mastocytosis and other hematological diseases. J. Leukoc. Biol. 67, 135–148

8. Porębska, N., Poźniak, M., Matynia, A., Żukowska, D., Zakrzewska, M., Otlewski, J., et al. (2021). Galectins as modulators of receptor tyrosine kinases signaling in health and disease. Cytokine Growth Factor Rev. 60, 89–106

9. Obata, Y., Toyoshima, S., Wakamatsu, E., Suzuki, S., Ogawa, S., Esumi, H., et al. (2014). Oncogenic Kit signals on endolysosomes and endoplasmic reticulum are essential for neoplastic mast cell proliferation. Nat. Commun. 5, 5715

10. Obata, Y., Horikawa, K., Takahashi, T., Akieda, Y., Tsujimoto, M., Fletcher, J. A., et al. (2017). Oncogenic signaling by Kit tyrosine kinase occurs selectively on the Golgi apparatus in gastrointestinal stromal tumors. Oncogene 36, 3661–3672

11. Obata, Y., Kurokawa, K., Tojima, T., Natsume, M., Shiina, I., Takahashi, T., et al. (2023). Golgi retention and oncogenic KIT signaling via PLCγ2-PKD2-PI4KIIIβ activation in gastrointestinal stromal tumor cells. Cell Rep. 42, 113035

12. Obata, Y., Hara, Y., Shiina, I., Murata, T., Tasaki, Y., Suzuki, K., et al. (2019). N822K-or V560G-mutated KIT activation occurs preferentially in lipid rafts of the Golgi apparatus in leukemia cells. Cell Commun. Signal. 17, 114

13. Yamawaki, K., Shiina, I., Murata, T., Tateyama, S., Maekawa, Y., Niwa, M., et al. (2021). FLT3-ITD transduces autonomous growth signals during its biosynthetic trafficking in acute myelogenous leukemia cells. Sci. Rep. 11, 22678

14. Hara, Y., Obata, Y., Horikawa, K., Tasaki, Y., Suzuki, K., Murata, T., et al. (2017). M-COPA suppresses endolysosomal Kit-Akt oncogenic signalling through inhibiting the secretory pathway in neoplastic mast cells. PLOS ONE 12, e0175514

15. Obata, Y., Horikawa, K., Shiina, I., Takahashi, T., Murata, T., Tasaki, Y., et al. (2018). Oncogenic Kit signalling on the Golgi is suppressed by blocking secretory trafficking with M-COPA in GISTs. Cancer Lett. 415, 1–10

16. Sztul, E., Chen, P. W., Casanova, J. E., Cherfils, J., Dacks, J. B., Lambright, D. G., et al. (2019). ARF GTPases and their GEFs and GAPs: concepts and challenges. Mol. Biol. Cell 30, 1249–1271

17. Casalou, C., Ferreira, A., and Barral, D. C. (2020). The Role of ARF Family Proteins and Their Regulators and Effectors in Cancer Progression: A Therapeutic Perspective. Front. Cell Dev. Biol. 8, 217

18. Takatsuki, A., and Tamura, G. (1985). Brefeldin A, a Specific Inhibitor of Intracellular Translocation of Vesicular Stomatitis Virus G Protein: Intracellular Accumulation of High-mannose Type G Protein and Inhibition of Its Cell Surface Expression. Agric. Biol. Chem. 49, 899–902

19. Lippincott-Schwartz, J., Yuan, L. C., Bonifacino, J. S., and Klausner, R. D. (1989). Rapid redistribution of Golgi proteins into the ER in cells treated with brefeldin A: evidence for membrane cycling from Golgi to ER. Cell 56, 801–813

20. Peyroche, A., Antonny, B., Robineau, S., Acker, J., Cherfils, J., and Jackson, C. L. (1999). Brefeldin A acts to stabilize an abortive ARF-GDP-Sec7 domain protein complex: involvement of specific residues of the Sec7 domain. Mol. Cell 3, 275–285

21. Sausville, E. A., Duncan, K. L., Senderowicz, A., Plowman, J., Randazzo, P. A., Kahn, R., et al. (1996). Antiproliferative effect in vitro and antitumor activity in vivo of brefeldin A. Cancer J. Sci. Am. 2, 52–58

22. Shiina, I., Umezaki, Y., Ohashi, Y., Yamazaki, Y., Dan, S., and Yamori, T. (2013). Total synthesis of AMF-26, an antitumor agent for inhibition of the Golgi system, targeting ADP-ribosylation factor 1. J. Med Chem. 56, 150–159

23. Shiina, I., Umezaki, Y., Murata, T., Suzuki, K., and Tonoi, T. (2018). Asymmetric total synthesis of (+)-coprophilin. Synthesis 50, 1301–1306

24. Watari, K., Nakamura, M., Fukunaga, Y., Furuno, A., Shibata, T., Kawahara, A., et al. (2012). The antitumor effect of a novel angiogenesis inhibitor (an octahydronaphthalene derivative) targeting both VEGF receptor and NF-κB pathway. Int. J. Cancer 131, 310–321

25. Rainbolt, T. K., Saunders, J. M., and Wiseman, R. L. (2014). Stress-responsive regulation of mitochondria through the ER unfolded protein response. Trends Endocrinol. Metab. 25, 528–537

26. Taguchi, T., Sonobe, H., Toyonaga, S., Yamasaki, I., Shuin, T., Takano, A., et al. (2002). Conventional and molecular cytogenetic characterization of a new human cell line, GIST-T1, established from gastrointestinal stromal tumor. Lab. Invest. 82, 663-665

27. Chung, B. M., Raja, S. M., Clubb, R. J., Tu, C., George, M., Band, V., et al. (2009). Aberrant trafficking of NSCLC-associated EGFR mutants through the endocytic recycling pathway promotes interaction with Src. BMC Cell Biol. 10, 84

28. Ménard, L., Floc’h, N., Martin, M. J., and Cross, D. A. E. (2018). Reactivation of Mutant-EGFR Degradation through Clathrin Inhibition Overcomes Resistance to EGFR Tyrosine Kinase Inhibitors. Cancer Res. 78, 3267–3279

29. Joffre, C., Barrow, R., Ménard, L., Calleja, V., Hart, I. R., and Kermorgant, S. (2011). A direct role for Met endocytosis in tumorigenesis. Nat. Cell Biol. 13, 827–837

30. Frazier, N. M., Brand, T., Gordan, J. D., Grandis, J., and Jura, N. (2018). Overexpression-mediated activation of MET in the Golgi promotes HER3/ERBB3 phosphorylation. Oncogene 38, 1936–1950

31. Slieker, L. J., Martensen, T. M., and Lane, M. D. (1986). Synthesis of epidermal growth factor receptor in human A431 cells. Glycosylation-dependent acquisition of ligand binding activity occurs post-translationally in the endoplasmic reticulum. J. Biol. Chem. 261, 15233–15241

32. Komada, M., Hatsuzawa, K., Shibamoto, S., Ito, F., Nakayama, K., and Kitamura, N. (1993). Proteolytic processing of the hepatocyte growth factor/scatter factor receptor by furin. FEBS Lett. 328, 25–29

33. Slee, E. A., Zhu, H., Chow, S. C., MacFarlane, M., Nicholson, D. W., and Cohen, G. M. (1996). Benzyloxycarbonyl-Val-Ala-Asp (OMe) fluoromethylketone (Z-VAD.FMK) inhibits apoptosis by blocking the processing of CPP32. Biochem J. 315, 21-24

34. Szul, T., Grabski, R., Lyons, S., Morohashi, Y., Shestopal, S., Lowe, M., et al. (2007). Dissecting the role of the ARF guanine nucleotide exchange factor GBF1 in Golgi biogenesis and protein trafficking. J. Cell Sci. 120, 3929–3940

35. Ignashkova, T. I., Gendarme, M., Peschk, K., Eggenweiler, H. M., Lindemann, R. K., and Reiling, J. H. (2017). Cell survival and protein secretion associated with Golgi integrity in response to Golgi stress-inducing agents. Traffic 18, 530–544

36. Shin, H. W., and Nakayama, K. (2004). Guanine nucleotide-exchange factors for arf GTPases: their diverse functions in membrane traffic. J. Biochem. 136, 761–767

37. Quilty, D., Gray, F., Summerfeldt, N., Cassel, D., and Melançon, P. (2014). Arf activation at the Golgi is modulated by feed-forward stimulation of the exchange factor GBF1. J. Cell Sci. 127, 354–364

38. Dell’Angelica, E. C., Puertollano, R., Mullins, C., Aguilar, R. C., Vargas, J. D., Hartnell, L. M., and Bonifacino, J. S. (2000). GGAs: a family of ADP ribosylation factor-binding proteins related to adaptors and associated with the Golgi complex. J. Cell Biol. 149, 81–94

39. Takatsu, H., Yoshino, K., Toda, K., and Nakayama, K. (2002). GGA proteins associate with Golgi membranes through interaction between their GGAH domains and ADP-ribosylation factors. Biochem. J. 365, 369–378

40. Dascher, C., and Balch, W. E. (1994). Dominant inhibitory mutants of ARF1 block endoplasmic reticulum to Golgi transport and trigger disassembly of the Golgi apparatus. J. Biol. Chem. 269, 1437–1448

41. Cohen, L. A., and Donaldson, J. G. (2010). Analysis of Arf GTP-binding protein function in cells. Curr. Protoc. Cell Biol. 1217, 1–17

42. Ip, C. K. M., Ng, P. K. S., Jeong, K. J., Shao, S. H., Ju, Z., Leonard, P. G., et al. (2018). Neomorphic PDGFRA extracellular domain driver mutations are resistant to PDGFRA targeted therapies. Nat. Commun. 9, 4583

43. Rieger, L., O’Shea, S., Godsmark, G., Stanicka, J., Kelly, G., and O’Connor, R. (2020). IGF-1 receptor activity in the Golgi of migratory cancer cells depends on adhesion-dependent phosphorylation of Tyr^1250^ and Tyr^1251^. *Sci. Signal.* 13, eaba3176

44. Schmidt-Arras, D., and Bo□hmer, F. D. (2020). Mislocalisation of Activated Receptor Tyrosine Kinases - Challenges for Cancer Therapy. Trends Mol. Med. 26, 833–847

45. Chiu, V. K., Bivona, T., Hach, A., Sajous, J. B., Silletti, J., Wiener, H., et al. (2001). Ras signalling on the endoplasmic reticulum and the Golgi. Nat. Cell Biol. 4, 343–350

46. Rocks, O., Peyker, A., Kahms, M., Verveer, P. J., Koerner, C., Lumbierres, M., et al. (2005). An acylation cycle regulates localization and activity of palmitoylated Ras isoforms. Science 307, 1746–1752

47. Pulvirenti, T., Giannotta, M., Capestrano, M., Capitani, M., Pisanu, A., Polishchuk, R. S., et al. (2008). A traffic-activated Golgi-based signalling circuit coordinates the secretory pathway. Nat. Cell Biol. 10, 912–922

48. Sato, I., Obata, Y., Kasahara, K., Nakayama, Y., Fukumoto, Y., Yamasaki, T., et al. (2009). Differential trafficking of Src, Lyn, Yes and Fyn is specified by the state of palmitoylation in the SH4domain. J. Cell Sci. 122, 965-975

49. Thomas, J. D., Zhang, Y. J., Wei, Y. H., Cho, J. H., Morris, L. E., Wang, H. Y., et al. (2014). Rab1A Is an mTORC1 Activator and a Colorectal Oncogene. Cancer Cell 30, 181–182

50. Bauer, S., Duensing, A., Demetri, G. D., and Fletcher, J. A. (2007). KIT oncogenic signaling mechanisms in imatinib-resistant gastrointestinal stromal tumor: PI3-kinase/AKT is a crucial survival pathway. Oncogene 26, 7560–7568

51. Green, D. R. (2022). Caspases and Their Substrates. Cold Spring Harb. Perspect. Biol. 14, a041012

52. Rokudai, S., Fujita, N., Hashimoto, Y., and Tsuruo, T. (2000). Cleavage and inactivation of antiapoptotic Akt/PKB by caspases during apoptosis. J. Cell. Physiol. 182, 290–296

53. Poreba, M., Strózyk, A., Salvesen, G. S., and Drag, M. (2013). Caspase substrates and inhibitors. Cold Spring Harb. Perspect. Biol. 5, a008680

54. Blay, J. Y., Kang, Y. K., Nishida, T., and von Mehren, M. (2021). Gastrointestinal stromal tumours. Nat. Rev. Dis. Primers 7, 22

55. Lasota, J., and Miettinen, M. (2008). Clinical significance of oncogenic *KIT* and *PDGFRA* mutations in gastrointestinal stromal tumours. Histopathology 53, 245–266

56. Schmitt, M. W., Loeb, L. A., and Salk, J. J. (2016). The influence of subclonal resistance mutations on targeted cancer therapy. Nat. Rev. Clin. Oncol. 13, 335–347

57. Mardones, G. A., Snyder, C. M., and Howell, K. E. (2006). Cis-Golgi matrix proteins move directly to endoplasmic reticulum exit sites by association with tubules. Mol. Biol. Cell 17, 525–538

58. Reiling, J. H., Olive, A. J., Sanyal, S., Carette, J. E., Brummelkamp, T. R., Ploegh, H. L., et al. (2013). A CREB3-ARF4 signalling pathway mediates the response to Golgi stress and susceptibility to pathogens. Nat. Cell Biol. 15, 1473–1485

59. Furitsu, T., Tsujimura, T., Tono, T., Ikeda, H., Kitayama, H., Koshimizu, U., et al. (1993). Identification of mutations in the coding sequence of the proto-oncogene c-kit in a human mast cell leukemia cell line causing ligand-independent activation of c-kit product. J. Clin. Invest. 92, 1736–1744

