## Supplementary Figures for "Brefeldin A and M-COPA block the export of RTKs from the endoplasmic reticulum via simultaneous inactivation of ARF1, ARF4, and ARF5"

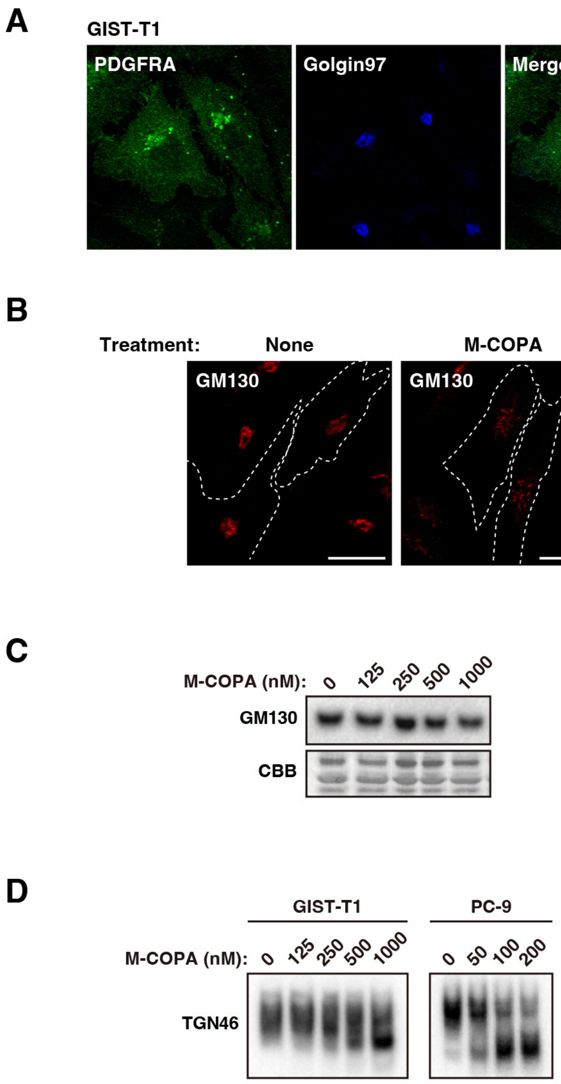


**Figure S1. M-COPA affects the distribution of GM130.**

*A,* GIST-T1 cells were immunostained for PDGFRA and golgin97. Bar, 20 μm.

*B* and *C,* GIST-T1 cells were treated with 1 μM M-COPA for 8 h. *B,* Cells were immunostained with Golgi matrix protein 130 kDa (GM130). Dashed lines indicate cell borders. Bars, 20 μm. *C,* Lysates were immunoblotted for GM130. Total protein levels were confirmed by Coomassie Brilliant Blue (CBB) staining.

*D,* GIST-T1 cells (left) or PC-9 cells (right) were treated with M-COPA for 8 h then immunoblotted for TGN46.

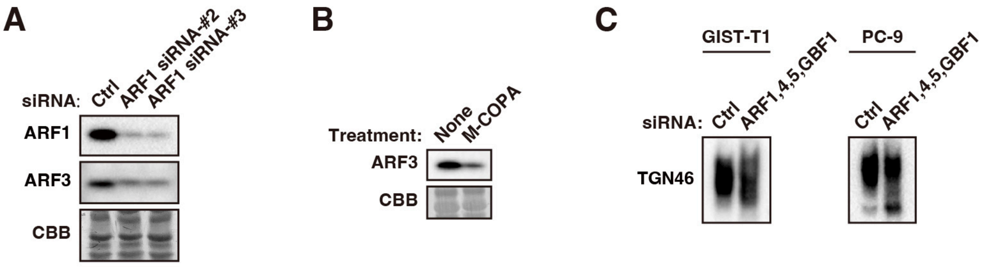


**Figure S2. ARF1 activity is involved in the maintain of ARF3 levels.**

*A,* GIST-T1 cells were transfected with the indicated siRNA for 48 h. Lysates were immunoblotted with the indicated antibodies. Total protein levels were confirmed by CBB staining.

*B,* GIST-T1 cells were treated with 1 $\mu$M M-COPA for 24 h and then immunoblotted for ARF3.

*C,* GIST-T1 cells (left) or PC-9 cells (right) were transfected with the indicated siRNAs for 48 h then immunoblotted for TGN46.

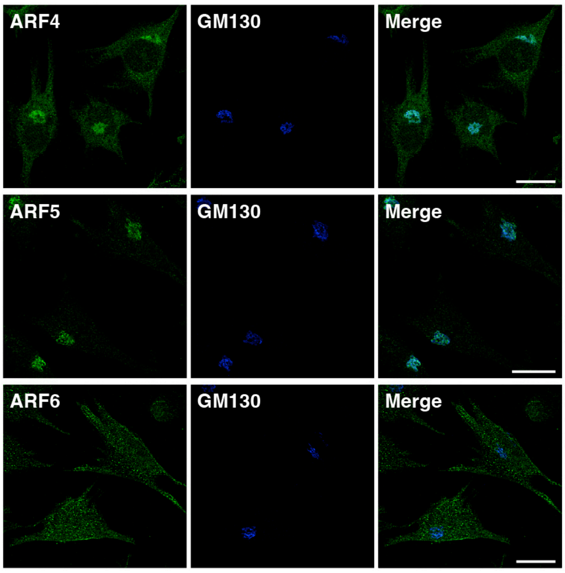


**Figure S3. ARFs are localized in the perinuclear region and the plasma membrane.**
GIST-T1 cells were immunostained for ARFs and Golgi matrix protein 130 kDa (GM130, Golgi marker, blue). Bars, 20 $\mu$m.

| Antibody | Host | Clone/catalog # | Distribution source |
|---|---|---|---|
| AKT | Mouse | 55 | BD Transduction Laboratories |
| AKT | Mouse | 40D4 | Cell Signaling Technology |
| AKT [pT308] | Rabbit | C31E5E | Cell Signaling Technology |
| ARF1 | Mouse | ARFS 1A9/5 | Santa Cruz Biotechnology |
| ARF1 | Rabbit | 10790-1-AP | Proteintech |
| ARF3 | Mouse | 41 | Santa Cruz Biotechnology |
| ARF3 | Mouse | 41 | BD Transduction Laboratories |
| ARF3 | Rabbit | 10800-1-AP | Proteintech |
| ARF4 | Rabbit | 11673-1-AP | Proteintech |
| ARF5 | Mouse | 14-07 | Santa Cruz Biotechnology |
| ARF5 | Rabbit | GTX104783 | GeneTex |
| ARF6 | Mouse | 3A-1 | Santa Cruz Biotechnology |
| ARF6 | Rabbit | 20225-1-AP | Proteintech |
| BIG1 | Mouse | G-3 | Santa Cruz Biotechnology |
| BIG2 | Mouse | H-6 | Santa Cruz Biotechnology |
| Cleaved caspase-3 | Rabbit | #9661 | Cell Signaling Technology |
| EGFR | Mouse | F4 | Santa Cruz Biotechnology |
| EGFR | Rabbit | D38B1 | Cell Signaling Technology |
| EGFR Δ746-750 | Rabbit | D6B6 | Cell Signaling Technology |
| EGFR [pY1068] | Rabbit | #2234 | Cell Signaling Technology |
| ERK1/2 | Rabbit | 137F5 | Cell Signaling Technology |
| ERK2 | Rabbit | K-23 | Santa Cruz Biotechnology |
| ERK [pT202/pY204] | Mouse | E10 | Cell Signaling Technology |
| ERK [pY204] | Mouse | E-4 | Santa Cruz Biotechnology |

**Supplementary Table 1.** List of primary antibodies.

| Antibody | Host | Clone/catalog # | Distribution source |
|---|---|---|---|
| GBF1 | Mouse | 25 | Santa Cruz Biotechnology |
| GBF1 | Mouse | 25 | BD Transduction Laboratories |
| GM130 | Mouse | 35 | BD Transduction Laboratories |
| GM130 | Rabbit | EP892Y | Abcam |
| Golgin97 | Mouse | CDF4 | Thermo Fisher Scientific |
| KIT | Mouse | E-1 | Santa Cruz Biotechnology |
| KIT | Mouse | 28 | BD Transduction Laboratories |
| KIT | Rabbit | D13A2 | Cell Signaling Technology |
| KIT | Rabbit | D3W6Y | Cell Signaling Technology |
| KIT [pY703] | Rabbit | D12E12 | Cell Signaling Technology |
| MET | Mouse | D-4 | Santa Cruz Biotechnology |
| MET | Rabbit | D1C2 | Cell Signaling Technology |
| MET [pY1234/1235] | Rabbit | D26 | Cell Signaling Technology |
| PDGFRA | Rabbit | D13C6 | Cell Signaling Technology |
| PDI | Mouse | RL-90 | Abcam |
| PERK | Mouse | B-5 | Santa Cruz Biotechnology |
| STAT3 | Mouse | 124H6 | Cell Signaling Technology |
| STAT3 [pY705] | Rabbit | D3A7 | Cell Signaling Technology |
| STAT5 | Mouse | 89 | BD Transduction Laboratories |
| STAT5 | Rabbit | D2O6Y | Cell Signaling Technology |
| STAT5 [pY694] | Rabbit | D47E7 | Cell Signaling Technology |

**Supplementary Table 2.** List of primary antibodies.

| Antibody | Host | Clone/catalog # | Distribution source |
|---|---|---|---|
| HRP anti-mouse IgG | Donkey | 715-035-151 | Jackson ImmunoResearch |
| HRP anti-rabbit IgG | Donkey | 711-035-152 | Jackson ImmunoResearch |
| AF488 anti-mouse IgG | Donkey | A21202 | Thermo Fisher Scientific |
| AF488 anti-rabbit IgG | Donkey | A21206 | Thermo Fisher Scientific |
| AF568 anti-mouse IgG | Donkey | A10037 | Thermo Fisher Scientific |
| AF568 anti-rabbit IgG | Donkey | A10042 | Thermo Fisher Scientific |
| AF647 anti-mouse IgG | Donkey | A31571 | Thermo Fisher Scientific |

**Supplementary Table 3.** List of secondary antibodies. HRP, horseradish peroxidase; AF, Alexa Fluor.

| Target | Catalog # | Distribution source |
|---|---|---|
| ARF1 | L-011580-00-0005 | Dharmacon |
| ARF1 | s1550 | Invitrogen |
| ARF1 | s1552 | Invitrogen |
| ARF3 | L-011581-00-0005 | Dharmacon |
| ARF4 | L-011582-00-0005 | Dharmacon |
| ARF5 | L-011584-00-0005 | Dharmacon |
| ARF6 | L-004008-00-0005 | Dharmacon |
| BIG1 | L-012207-00-0005 | Dharmacon |
| BIG2 | L-012208-02-0005 | Dharmacon |
| GBF1 | L-019783-00-0005 | Dharmacon |
| Non-targeting control siRNAs | D-001810-10-20 | Dharmacon |
| Negative control siRNA | 4390843 | Invitrogen |

**Supplementary Table 4.** List of siRNAs.
